# A parameter-efficient deep learning approach to predict conversion from mild cognitive impairment to Alzheimer’s disease

**DOI:** 10.1101/383687

**Authors:** Simeon Spasov, Luca Passamonti, Andrea Duggento, Pietro Liò, Nicola Toschi

**Affiliations:** University of Cambridge, Cambridge, Department of Computer Science and Technology, William Gates Building, 15 J J Thomson Ave, Cambridge, CB3 0FD, UK.; Department of Clinical Neurosciences, University of Cambridge, Herchel Smith Building, Forvie Site, Robinson Way, Cambridge Biomedical Campus, Cambridge CB2 0SZ Cambridge.; Department of Biomedicine and Prevention, University of Rome “Tor Vergata”, Via Cracovia, 00133 Roma RM, Italy.; A.A. Martinos Center for Biomedical Imaging, Massachusetts General Hospital and Harvard Medical School, Boston (USA).

**Author notes:** these authors contributed equally to this publication (Pietro Liò, Nicola Toschi).

**Keywords:** deep learning, neural networks, classification, Mild Cognitive Impairment, Alzheimer’s disease, Magnetic resonance imaging, ADNI, Early diagnosis

## Abstract

Some forms of mild cognitive impairment (MCI) can be the clinical precursor of severe dementia like Alzheimer’s disease (AD), while other types of MCI tend to remain stable over-time and do not progress to AD pathology. To choose an effective and personalized treatment for AD, we need to identify which MCI patients are at risk of developing AD and which are not.

Here, we present a novel deep learning architecture, based on dual learning and an ad hoc layer for 3D separable convolutions, which aims at identifying those people with MCI who have a high likelihood of developing AD. Our deep learning procedures combine structural magnetic resonance imaging (MRI), demographic, neuropsychological, and APOe4 genotyping data as input measures. The most novel characteristics of our machine learning model compared to previous ones are as follows: 1) multi-tasking, in the sense that our deep learning model jointly learns to simultaneously predict both MCI to AD conversion, and AD vs healthy classification which facilitates the relevant feature extraction for prognostication; 2) the neural network classifier employs relatively few parameters compared to other deep learning architectures (we use ~550,000 network parameters, orders of magnitude lower than other network designs) without compromising network complexity and hence significantly limits data-overfitting; 3) both structural MRI images and warp field characteristics, which quantify the amount of volumetric change compared to the common template, were used as separate input streams to extract as much information as possible from the MRI data. All the analyses were performed on a subset of the Alzheimer’s Disease Neuroimaging Initiative (ADNI) database, for a total of n=785 participants (192 AD, 409 MCI, and184 healthy controls (HC)).

We found that the most predictive combination of inputs included the structural MRI images and the demographic, neuropsychological, and APOe4 data, while the warp field metric added little predictive value. We achieved an area under the ROC curve (AUC) of 0.925 with a 10-fold cross-validated accuracy of 86%, a sensitivity of 87.5% and specificity of 85% in classifying MCI patients who developed AD in three years’ time from those individuals showing stable MCI over the same time-period. To the best of our knowledge, this is the highest performance reported on a test set achieved in the literature using similar data. The same network provided an AUC of 1 and 100% accuracy, sensitivity and specificity when classifying NC from AD. We also demonstrated that our classification framework was robust to different co-registration templates and possibly irrelevant features / image sections.

Our approach is flexible and can in principle integrate other imaging modalities, such as PET, and a more diverse group of clinical data. The convolutional framework is potentially applicable to any 3D image dataset and gives the flexibility to design a computer-aided diagnosis system targeting the prediction of any medical condition utilizing multi-modal imaging and tabular clinical data.

## Introduction

More than 30 million people have a clinical diagnosis of Alzheimer’s disease (AD) worldwide, and this number is expected to triple by 2050 (Barnes and Yaffe, 2011), also due to increased life expectancy and improvements in care (Ferri et al., 2005). AD is a form of dementia characterized by extracellular β-amyloid peptide plaque deposits and abnormal tau accumulation and phosphorylation which ultimately lead to neuronal and synaptic loss (Murphy et al. 2010). AD-related neurodegeneration follows specific patterns which arise from subcortical areas and spread to the cortical mantle (Braak and Braak et al. 1996). The classic clinical hallmark of the most common form of AD (i.e., the amnestic type) are impairments in episodic memory, followed by visuo-spatial and orientation problems, and ultimately by frank dementia.

Mild cognitive impairment (MCI) is a wide and heterogeneous spectrum of disorders which causes relatively less acute and noticeable memory deficit than AD. Around 10%-15% of MCI patients per year convert to AD over a short observation period (Braak and Braak, 1995; Mitchell and Shiri-Feshki, 2008), although the annual conversion rate diminishes with time to form a mean annual conversion rate of ~4%. MCI patients who do not convert to AD tend to either remain stable, develop other forms of dementia, or even revert to a healthy state, which suggests that MCI is a heterogeneous combination of disorders which are likely to be associated with several distinct etio-pathogenetic mechanisms. In this context, AD-related neuropathological markers have been observed several years before clinical manifestation of memory symptoms (Braak and Braak, 1996; Delacourte et al., 1999; Morris et al., 1996; Serrano-Pozo et al., 2011; Mosconi et al., 2007), which suggests that AD development could be predicted before clinical onset via in vivo biomarker analysis (e.g. PET and MR imaging as well as blood or cerebrospinal fluid (CSF) biomarkers) (Markesbery, 2010; Baldacci et al., 2018; Hampel et al. 2018; Teipel et al., 2018). Magnetic resonance imaging (MRI) has garnered interest in AD diagnosis as well as prediction of MCI to AD conversion. Relative to cerebrospinal fluid (CSF) and positron emission tomography (PET) biomarkers, MRI measures have the notable advantages of not using ionizing radiation, of being noninvasive, less expensive and more widely available in less specialized medical environments. MRI markers also enable the possibility to gather multimodal information (e.g. structural and functional) within the same scanning session.

For these reasons, there has been a growing interest in developing MRI-based computational tools to discriminate AD patients from healthy individuals, as well as (most importantly) in distinguishing between stable MCI (sMCI) patients and MCI patients who progress (pMCI) to AD. To this end, different clinical data and imaging modalities have been employed with variable rates of success, including PET studies (Choi et al. 2018; Mosconi et al. 2004, Mosconi et al. 2007, Shaffer et al. 2013, Young et al. 2013), MRI studies (Filipovych et al. 2011; Moradi et al. 2015; Mosconi et al. 2007; Tong et al. 2017, Young et al. 2013), cognitive testing studies (Casanova et al. 2011; Moradi et al. 2015), and CSF biomarker studies (Davatzikos et al. 2011; Hansson et al. 2006; Riemenschneider et al. 2002; Sonnen et al. 2010). As an example, Moradi et al. 2015 as well as Tong et al. 2017 first perform feature selection to extract informative voxels from MRI volumes via regularized logistic regression, and subsequently use the extracted voxels, along with cognitive measures, to produce support vector machine (SVM)-based predictions, achieving an area under the Receiver Operating Characteristic (ROC) curve (AUC) between 0.9 and 0.92. In the case of Hojjati et al., 2017, who use baseline resting state fMRI data and achieve an AUC of 0.95, features are engineered by constructing a brain connectivity matrix which is treated as a graph, and the extracted graph measures are inputted into a SVM.

Most of the above-mentioned studies employ a classification pipeline which relies on two independent steps. First, a dimensionality reduction method, such as ICA (Shaffer et al. 2013), L1 regularization (Moradi et al. 2015; Tong et al. 2017) or morphometry (Davatzikos et al. 2011; Fan et al. 2007), is used to reduce the raw images or volumes to a relatively small number of (hopefully) highly descriptive factors. Then, these factors are fed into a multivariate pattern classification algorithm. Notably, the dimensionality reduction and classification algorithms are two separate mathematical models which involve different assumptions, hence possibly resulting in loss of relevant information in the classification process (Nguyen and Torre, 2010). Also, the most frequently used classifiers, such as SVM (Moradi et al., 2015; Hojjati et al., 2017, Tong et al., 2017) and Gaussian Processes (Young et al., 2013), require the use of kernels, or data transformations, chosen from a limited user-specified set, which map the data to a new space in the hope that it will be more easily separable. However, constructing or choosing an application-specific kernel to act as a reasonable similarity measure for the task at hand is not always possible.

The use of two disjoint pipelines and the need to construct ad-hoc kernels can be surmounted by the use of a class of algorithms known as deep learning, which afford much greater representational flexibility than kernel-based methods and also automatically “learn” data transformations which maximize an arbitrary performance metric. Such methods have been applied to AD vs. healthy subject discrimination (Hosseini-Asl et al., 2016; Liu et al., 2018; Payan and Montana, 2015) and pMCI vs sMCI classification (Choi et al., 2018; Lu et al., 2018a, b) As an example, Choi et al., 2018 and Lu et al., 2018a use deep learning to achieve one of the highest pMCI/sMCI classification performances to-date ( ~84% - 82% conversion rate accuracies for these studies respectively). Their predictions are based on a single (albeit very informative) imaging modality (PET) which employs ionizing radiation. A comparison between recent studies and methods is provided in Table 3.

As is well known, the superior representational capacity of deep learning methods relies on a high number of neural network parameters. Frequently, this gives rise to overfitting, i.e. a satisfactory training performance which however does not generalize well to unseen samples during testing or when applying the model. Although it has been demonstrated that deep learning approaches can yield impressive performance, the data-scarce nature of medical datasets is not commonly sufficient to build a useful network architecture. The aim of this paper is therefore to develop and employ a parameter-efficient neural network architecture, based on more recent convolutional neural network layers, i.e. namely 3D separable and grouped convolutions, which were developed specifically for computer vision tasks. Additionally, we implement a joint/dual-learning approach which simultaneously learns multi-task classification of pMCI vs. sMCI and AD vs Health Controls (HC) and combines several input streams (including structural MRI as well as clinical variables comprising demographic, neuropsychological, and APOe4 genotyping data). These newer network designs have been shown to yield superior performance on generic visual discrimination problems like ImageNet (Russakovsky et al., 2015; Chollet et al., 2016) while maintaining the overall network parameter count low, hence efficiently battling the overfitting problem. Additionally, we developed a novel feature extractor sub-network and, in order to employ these methods efficiently, we combined the Tensorflow (Abadi et al., 2016) and Keras (Chollet et al., 2015) libraries with our own implementation of 3D separable convolutions (code available freely upon request).

## Methods

### 1. Participants and data

All data was obtained from the Alzheimer’s Disease Neuroimaging Initiative (ADNI) and comprised 435 men and 350 women aged between 55 and 91 years. The majority of subjects identified as white (>94%) and non-Hispanic (99.98%). All data we used is summarized in Table 1. Differences in median age across groups were tested using Friedman’s ANOVA and group x gender interactions were tested using Fisher’s exact test. None of these interactions resulted statistically significant (p> 0.05). For all participants, we employed the Magnetization Prepared Rapid Gradient-Echo (MPRAGE) T1-weighted image (structural MRI) as well as the following data: demographic data (age, gender, ethnic and racial categories, education), neuropsychological cognitive assessment tests like the dementia rating scale (CDRSB), the Alzheimer’s disease assessment scale (ADAS11, ADAS13), episodic memory evaluations in the Rey Auditory Verbal Learning Test (RAVLT), as well as APOe4 genotyping. All of the data we use in this study is from baseline assessments (no longitudinal data is used).

**Table 1.** Demographic, neuropsychological and cognitive assessment as well as APOe4 genotyping data used in this study. The data is presented in a mean±std format. Abbreviations: APOe4 - Apolipoprotein E; CDRSB – Clinical Dementia Rating Sum of Boxes; ADAS – Alzheimer’s Disease Assessment Scale; RAVLT – Ray Auditory Verbal Learning Test.

|  | No. of subjects | Age (years) | Male/Female | years in education | APOe4 expression |  |  | CDRSB | ADAS11 | ADAS13 | RAVLT |  |  |  |
| --- | --- | --- | --- | --- | --- | --- | --- | --- | --- | --- | --- | --- | --- | --- |
|  |  |  |  |  | 0 | 1 | 2 |  |  |  | immediate | learning | forgetting | % forget |
| AD | 192 | 75.6±7 | 103/81 | 15±2.9 | 57 | 86 | 41 | 4.4±1.6 | 18.8±6 | 29±7.3 | 23±7 | 1.7±1.8 | 4.4±1.9 | 89.4±21.2 |
| HC | 184 | 74.6±6 | 92/100 | 16.3±2.7 | 144 | 43 | 5 | 0.2±0.9 | 6±3.8 | 9.3±5.7 | 44±10.5 | 6±2.4 | 3.7±2.7 | 33.1±27.7 |
| pMCI | 181 | 73.7±7 | 108/73 | 15.9±2.8 | 61 | 90 | 30 | 2±1 | 13.5±4.2 | 21.9±5.5 | 27.2±6.5 | 2.9±2.2 | 4.9±2.1 | 78.3±27 |
| sMCI | 228 | 72.2±7 | 132/96 | 16±2.8 | 145 | 67 | 16 | 1.2±0.6 | 8.4±3.3 | 13.5±5.3 | 38.5±10 | 4.75±2.5 | 4.35±2.6 | 50±30 |

### 2. Data Preprocessing

Prior to classification, all T1 weighted (T1w) images were registered to a common space (i.e. T1 template). In detail, two different T1 templates were used in order to assess the robustness of our classification methodology to coregistration inaccuracies. First, we built a custom T1 template specific to this study. To this end, we employed all T1w images, which (after N4 bias field correction) where nonlinearly co-registered to each other and averaged iteratively (i.e. the group average was recreated at the end of each iteration). The procedure was based on symmetrical diffeomorphic mapping and employed five total iterations. The second template was the Montreal Neurological T1 Template (MNI152_T1_1mm). After the creation of both templates, all single-subject T1w images were nonlinearly registered to both templates. After co-registration to both templates we also extracted the local Jacobian Determinant (JD) images of the nonlinear part of the deformational field taking each image into template space, and masked out all non-brain areas using brainmasks generated in template space using BET, part of FSL (Jenkinson et al., 2012). The JD maps were used to complement the MRI images as an additional input stream in our model (see below). Additionally, in order to evaluate how much a priori knowledge about AD brain pathophsyiology could improve our classification and also how much irrelevant features hamper classification performance, we defined a set of regions of interest (ROIs) which included only brain areas known to be heavily involved in AD-related atrophy, namely parietal, temporal and frontal lobes in order to perform an inclusion test (see fig. 5). This was based on the Hammers et al. 2003 atlas^© Copyright Imperial College of Science, Technology and Medicine 2007 (www.brain-development.org)^.

All template creation and registration procedures were performed using the ANTs package (Avants et al., 2010, Avants et al., 2011). In detail, the high-dimensional non-linear transformation (symmetric diffeomorphic normalization transformation) model was initialized through a generic linear transformation which consisted of center of mass alignment, rigid, similarity and fully affine transformations followed by (metric: neighbourhood cross correlation, sampling: regular, gradient step size: 0.12, four multi-resolution levels, smoothing sigmas: 3, 2, 1, 0 voxels in the reference image space, shrink factors: 6, 4, 2, 1 voxels. We also used histogram matching of images before registration and data winsorisation with quantiles: 0.001, 0.999. The convergence criterion was set to be as follows: slope of the normalized energy profile over the last 10 iterations < 10-8). Co-registration of all scans required approximately 19200 hour of CPU time on a high- performance parallel computing cluster.

Numerical normalization for the co-registered MRI images was performed per sample, i.e. each 3D volume was standardized to 0 mean and unit standard deviation. The reasoning behind this is that brain atrophy could be recognized as an in-sample shift in intensity for a certain area compared to other regions. The normalization applied to the clinical features, i.e. the demographic, neuropsychological, and APOe4 genotyping data, also follows the same feature scaling procedure, where the values of each separate clinical factor are normalized between [0, 1]. On the other hand, the extracted JD images were feature-scaled to have voxel values in the [0;1] range via subtracting the smallest value in the entire JD image set, and dividing by the difference between the largest and smallest values (also in the entire JD image set). This retains class-wise differences in volumetric changes created when co-registering an image to a template while rescaling the data to a global maximum and minimum.

### 3. Deep Learning Architecture

#### 3.1. Architecture Overview

A high-level overview of the network design is shown in fig. 2. In this paper, we developed a feature extractor sub-network (referred to as the *multi-modal feature extractor* in fig. 2), inspired by the parameter-efficient separable and grouped convolutional layers presented in AlexNet (Krizhevsky et al., 2012) and Xception (Chollet, 2017, Velickovic et al., 2016). In detail, the layers of the feature extractor are shared between two tasks - MCI-to-AD conversion prediction and AD/HC classification (see fig. 3 and fig. 4). The assumption is that both problems share common underlying factors, i.e. the MCI subjects who convert lie on a continuum between HC and AD. This means similar data transformations are likely to be useful for prediction of the two different problems. Also, this procedure increases the number of samples the extractor network is trained on, hence reducing overfitting. Also, balancing between the tasks can be seen as imposing soft constraints on the network parameters, and if some of the factors that explain the variations in our data are shared between the two discrimination problems, overfitting is reduced further. The feature extractor sub-network extracts 4-dimensional vectors for each of the two classification problems. These resulting latent representations are then processed by two separate fully connected layers (see fig. 1) with sigmoid activations and a binary cross-entropy loss applied at the output of each. The outputs of the fully connected layers are in the 0 to 1 range. The closer the activation is to 1, the more confident the model is that the input pattern corresponds to a diseased individual (i.e. AD or pMCI, depending on the classification task), and vice versa.

**Fig. 1.**
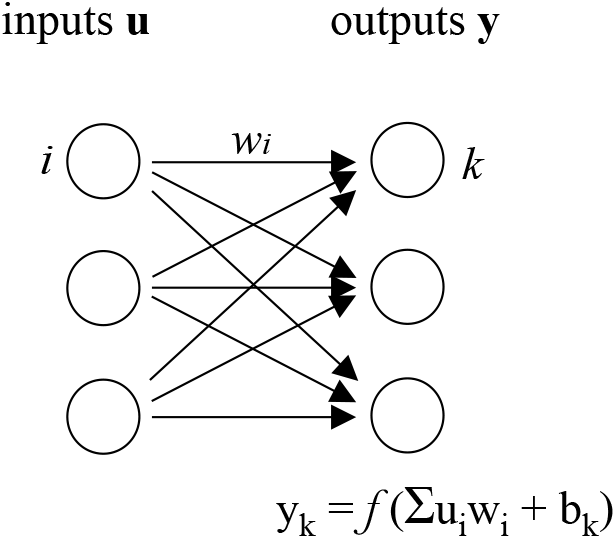
Operation of a dense or fully connected layer. The outputs y_k_ are formed as a non-linear transformation of the input vector **u**. The non-linear activation works on a weighted sum of the inputs, 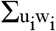, and a bias term b_k_. These layers are employed to process the clinical inputs in the Multi-modal feature extractor and to produce the output labels of our model.

**Fig. 2.**
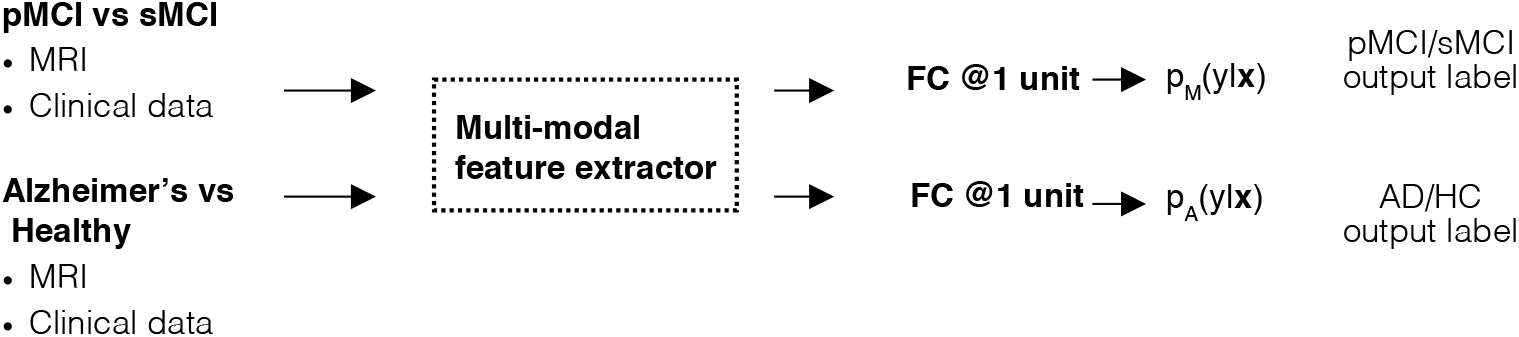
Overview of our multi-tasking neural network methodology. We have designed a sub-network (the multi-modal feature extractor) to extract 4-d feature representations from the inputs of both tasks/datasets. This sub-network (with **θ** network parameters) is applied on the data from both the pMCI/sMCI and AD vs healthy discrimination problems, as we assume the underlying factors of the conditions are similar, hence similar data transformations are likely to be useful. We then employ two fully connected layers, parametrized by **φ** and **ψ**, with sigmoid outputs. The sigmoid outputs approximate the conditional distribution of the labels for the two problems given the inputs (p_A_(y|**x**) for the AD vs healthy task and p_M_(y|**x**) for the pMCI vs sMCI task). We learn the network parameters such that our model outputs correspond to the true labels in the dataset by minimizing the binary cross-entropy between the observed and estimated targets. The multi-modal feature extractor is represented by a dashed-line rectangle in fig. 2 and fig. 4.

**Fig. 3.**
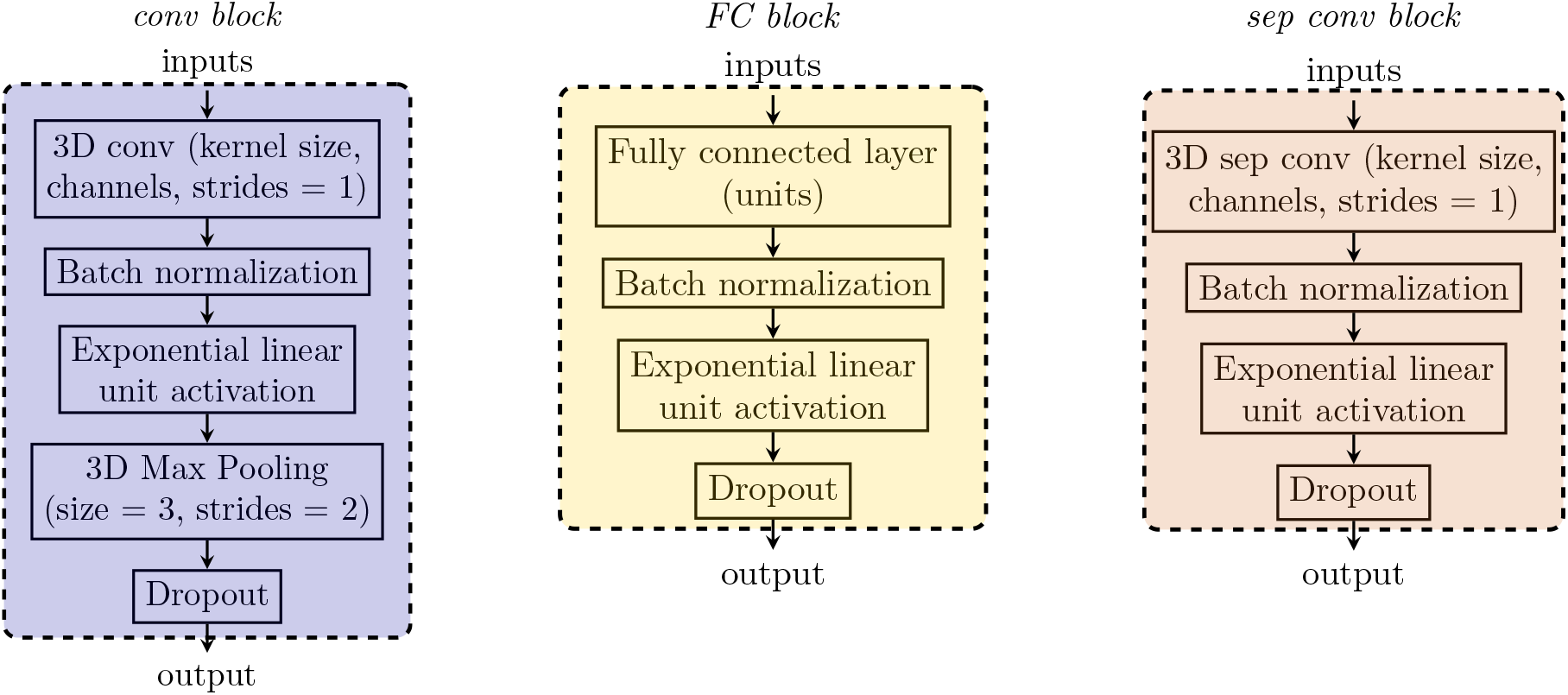
Implementation of the convolutional, fully connected and separable convolutional blocks (conv block, FC and sep conv block respectively). These blocks comprise several sequential operations – firstly a (separable) convolutions or dense layer followed by batch normalization and an ELU activation function. Conv blocks utilize 3D Max pooling with a window size of 3 and strides of 2 to gradually decrease input image dimensionality. Dropout is applied in all operational blocks. Convolutional, fully connected and max pooling layer require us to define hyperparameters, such as kernel size, number of units, etc. These are given in brackets with some commonly used default values for our network design.

**Fig. 4.**
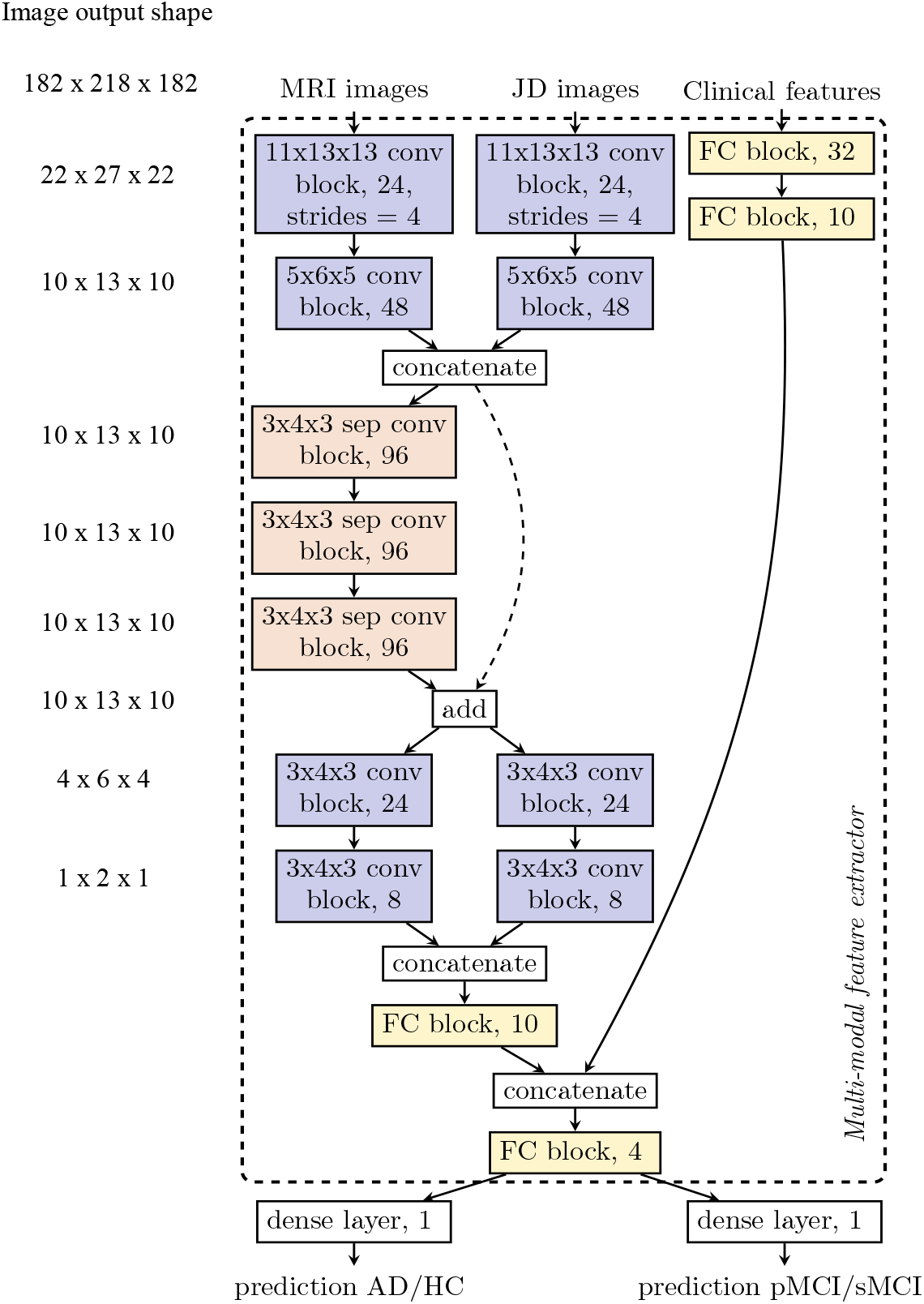
The architecture of the neural network designed to take multiple 3D image volumes and tabular clinical inputs. The design of the network relies on the operational blocks shown in fig. 3. For conv and sep conv blocks we use the notation: *kernel size, (sep) conv block, output channels*. If the strides are different from the default value of 1, the new stride value is shown in addition at the end. The *concatenation* operation works by merging the activation maps along the channel axis. Addition in the *add* block is performed element-wise between two sets of activation maps of the same size along all dimensions. The operational blocks are color-coded for the ease of the reader both in fig. 3 and fig. 4. Our network relies on decreasing the dimensionality of the image inputs using standard, separable and grouped convolutional blocks before concatenating the image embeddings with the compressed via fully connected blocks clinical features. The separable and grouped convolutions allow us to process the images in a parameter-efficient manner while the residual connection (dashed arrow from *concatenate* to *add*) facilitates training (Chollet, 2017) The multi-modal feature extractor sub-network (within the dashed rectangle) outputs 4-d embeddings of the input data and passes it to a dense layer which produces a prediction score. The same multi-modal feature extractor processes the inputs from both the MCI/HC and pMCI/sMCI tasks. Two different dense layers produce the final prediction scores for the two classification problems, however.

#### 3.2. Mathematical formulation of Model

We will denote the input data and labels as pairs (**X, Y**) = {(**x**^A^_1_, y^A^_1_),…, (**x**^A^_N_, y^A^_N_),…, (**x**^M^_1_, y^M^_1_),…, (**x**^A^_N’_, y^A^_N’_)}, where **x**^A^_i_ is the i-th observation from the Alzheimer’s and healthy subset, and **x**^M^_j_ is the j-th observation from the pMCI vs sMCI subset. Both classification problems have corresponding class labels y^A^_i_ and y^M^_j_ ∈ {0, 1}. We refer to the empirical distributions over the AD/HC and MCI subsets as 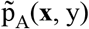 and 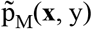 respectively. The model log likelihoods (i.e. the conditional probabilities of the target variables, y, given the input data **x** which we model with the neural network) for the two classification problems are given by:

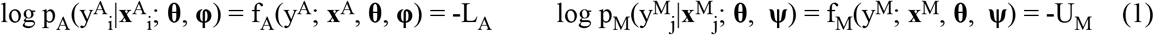

The likelihood functions f_A_ and f_M_ are modelled as Bernoulli distributions, parametrized by neural network-based transformations of the input data as described in fig. 2 The goal is to learn the network parameters such that we can approximate the *true* conditional probabilities of the labels given the inputs via the likelihood functions given by eq. 1. We use **θ** to denote the parameters in the multi-modal feature extractor sub-network, and **φ** and **ψ** to denote the weights in the final fully connected layers that output the class probabilities for the Alzheimer’s vs healthy and pMCI vs sMCI tasks respectively. Learning the network parameters can be represented as:

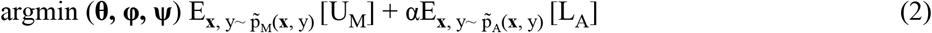

As U_M_ and L_A_ represent negative log-likelihoods, the objective function given in eq. (2) can be viewed as minimizing the weighted sum between two binary cross-entropy terms between the observed and estimated (by our network) class probabilities. Intuitively, learning the network parameters is akin to maximizing the probability of observing the labels in both datasets under the model, given the input cognitive, genetic and MRI biomarkers. We also introduced the α hyperparameter to control the trade-off between the two tasks during learning, and use α = 0.25 in all experiments. This is a heuristic choice based on the observation that the AD/HC problem is much easier than the pMCI/sMCI problem and that the model quickly achieves high validation accuracy when α = 0.25.

#### 3.3. 3D Convolutions

Convolutional layers employed in our study work by convolving an input tensor, **x**, with a kernel of weights **W**, then adding a bias term *b*, and finally passing the result through a non-linearity. To extract a rich set of representations we repeat this process with K different kernels (also known as channels or filters) convolving the same tensor **x**, each resulting in a new *feature map* **h_k_**. Hence, we can write:

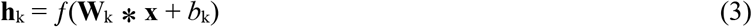

The feature map subscript is k = [1,…, K]. The function *f* can be selected from a range of differentiable non-linear transformations, such as the sigmoid f(u) = (1 + exp(−u))^−1^ and the exponential linear unit, or ELU, (Clevert et al. 2015): f(u) = u if u >= 0 and f(u) = exp(u) - 1 if u<0. We employ the ELU transformation in our hidden layer activations and a sigmoid output for label predictions. The set of K feature maps extracted from the input **x** defines a single layer ℓ = [1,…, L] in our convolutional neural network. Thus, the k^th^ feature map at layer ℓ is denoted as **h**_k_^ℓ^. To construct a hierarchy of features we can use the outputs of layer ℓ-1 as inputs to layer ℓ:

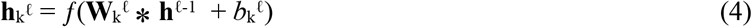

where **h**^0^ is **x**. Note that in eq. (2), **h**^ℓ-1^ = [**h**_0_^ℓ-1^,…, **h**_K_^ℓ-1^] is a 4-D tensor - a collection of the K 3D feature maps extracted at layer ℓ-1. Consequently, **W**_k_^ℓ^ is also a 4-D tensor kernel of size N^1^xN^2^xN^3^xK. This filter is multiplied element-wise during convolution with a N^1^xN^2^xN^3^ patch in each of the K feature maps and the result is summed to produce a single scalar element (after adding a bias term and passing through a non-linear function). The convolutional procedure can be seen as sliding this kernel with strides in all three dimensions to produce **h**_k_^ℓ^. It is important to note that the number of parameters needed to extract K^ℓ^ feature maps in layer ℓ from the K^ℓ-1^ feature maps in layer ℓ-1 is given by:

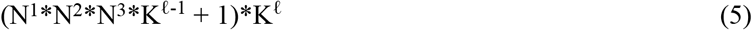

where N^1^xN^2^xN^3^ is the filter size used (see section 3.8 for actual values used in this paper).

#### 3.4. Fully connected (Dense) Layers

Fully connected (FC) layers are designed to work on vectorized inputs u. The operation of the dense layer is depicted in fig. 1. Each input u_i_ has an associated weight w_i_. In order to produce an output y_k_, we form the weighted sum of all inputs 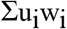, then add a bias term b_k_, and pass the result through a differentiable non-linear function like the sigmoid or the exponential linear unit. We can repeat this procedure K times with different weight parameters to produce an output vector **y**, which can be used as an input to another fully connected layer. In our work we employ these dense connections to process the tabular clinical features and to produce the final output predictions (or probability scores) of our model.

#### 3.5. Batch normalization, dropout, L2 regularization

Several strategies are used in our network to battle overfitting. The first one is batch normalization (Ioffe and Szegedy 2015) which normalizes a layer’s outputs by subtracting their mean and dividing by their standard deviation. This whitening procedure enforces a fixed distribution of activations which stabilizes and accelerates the rate of training of deep neural nets. We also implement dropout (Srivastava et al. 2014), which works by randomly dropping units and their connections during training. An intuitive explanation of its efficacy is that each unit must learn to extract useful features on its own with different sets of randomly chosen inputs. As a result, each hidden unit is more robust to random fluctuations and learns a generally useful transformation. Finally, L2 regularization penalizes weights of high absolute value, hence directly limiting the capacity of our model, i.e. improving overfitting.

#### 3.6. Separable Convolutions

The separable convolutions we employ are similar to standard convolutional layers but reformulate the procedure in two steps by performing *depthwise* and then *pointwise* operations. Firstly, each input channel is spatially convolved separately, then the resulting outputs are mixed via *pointwise* convolutions with a kernel size of 1 × 1 × 1. The depthwise procedure simply reformulates the convolutional operation from eq. (4) to:

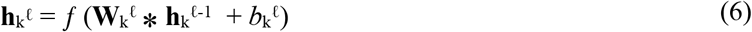

Note that the difference between eq. (4) and eq. (6) is the subscript k in **h**_k_^ℓ-1^, denoting that feature map k in layer ℓ (**h**_k_^ℓ^) is only a function of feature map k in layer ℓ-1 (**h**_k_^ℓ-1^) in the separable convolutions case. On the other hand, standard convolutions take as an input all K^ℓ-1^ feature maps to produce a single output. Consequently, with our approach the parameter count in **W**_k_^ℓ^ is reduced to (N1*N2*N3+1)*K^ℓ^, which is ~K^ℓ-1^ times more parameter-efficient as compared to standard convolutions (eq. (5)). The pointwise operation mixes all channels and requires K^ℓ^*K^ℓ-1^ parameters. Hence, the overall number of weights in separable convolutions is given by:

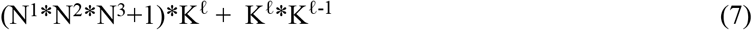

Considering the kernel sizes and number of filters in our network architecture, substituting a single conventional convolutional layer with a separable one results in ~20 times less parameters for that layer. In order to achieve the above operations, we implemented an ad-hoc 3D separable convolution module as a custom Keras layer based on a TensorFlow backend.

#### 3.7. Grouped Convolutions

The grouped layer can be viewed as a compromise between standard convolutions and the separable case. This procedure splits the previous layer’s feature maps in two groups (G1 and G2) along the channel axis and treats them as separate when applying further transformations (see fig. 4). As a result, only half of the channels are used to produce a single output feature map. The grouped layer requires twice fewer parameters than the standard convolutional approach, assuming the same overall number of output feature maps is generated.

#### 3.8. Network architecture

Since several different sequences of layers are frequently reused, they are combined in operational blocks. Each block follows a similar pattern. For instance, convolutional blocks, or conv blocks, used to processes the 3D MRI tensors, comprise a convolutional kernel with linear activations, batch normalization and an exponential linear unit (ELU) transformation with dropout. In order to reduce the resulting spatial dimensions, max pooling is used, where only the highest value in an image patch is retained, with a window of 3 pixels and a stride of 2. Each operation is applied to the outputs of the previous one. On the other hand, the clinical features undergo a series of transformations by dense, or FC (fully connected), blocks. Since these blocks act on vectorized inputs, a linear dense layer is employed instead but the same regularization precautions and activations as in the conv block are applied. We also implement a separable convolutions block, or sep conv block, which resembles the conv block but substitutes traditional convolutions with separable ones and does not rely on any pooling operations. All of these blocks are depicted in fig. 3. Fig. 4 shows the neural network architecture we use for the AD/HC and pMCI/sMCI classification problems. Firstly, two consecutive convolutional blocks are used to reduce the dimensionality of the input MRI and Jacobian images. We then concatenate the outputs of the second conv block from the MRI and Jacobian images along the channel axis. The majority of the feature extraction is then performed by three sequential separable convolutional blocks. The dimensionality of the activation maps remains the same during this procedure. The output from the last sep conv block is summed element-wise with the activation maps from the second conv block (also known as a residual connection, introduced in He et al., 2015 and Chollet, 2017. It has been shown that residual connections facilitate training as the depth of the neural network increases. We now split the result of the summation along the channel axis in two groups to perform a grouped convolution. The motivation behind opting for grouped convolutions is to further reduce the dimensionality of the activation maps which is not possible by using the fully separable convolutions as outlined in eq. 6 but is more parameter-efficient than utilizing traditional convolutions. At this stage of the image processing pipeline the shape of the activation maps is 1 × 2 × 1 with 16 channels after concatenation (8 channels in each group). We flatten the feature maps to a 32-dimensional vector and apply a fully connected block with 10 output units. This 10-dimensional vector forms the final embedding of the MRI and Jacobian images. The clinical features undergo 2 sequential transformations by fully connected blocks with 32 and 10 units respectively. The clinical features and image embeddings are concatenated and processed by a fully connected block with 4 output units. All of these operations acting on the MRI, Jacobian and clinical feature inputs which ultimately compress the input data in a 4-dimensional vector comprise the *Multi-modal feature extractor*. The parameters associated with the multi-modal feature extractor are denoted by **θ** in the mathematical formulation of our model in section 3.2. In order to obtain a prediction for each of the two tasks (AD/HC and pMCI/sMCI) we pass the 4-d output of the feature extractor sub-network through two dense (fully connected) *layers* (not blocks) with sigmoid activations and single output units. We use **φ** and **ψ** in our mathematical formulation to denote the weights in these final fully connected layers which model the class probabilities for the AD/HC and pMCI/sMCI tasks respectively.

The overall parameter count of our neural network model is 557,000. Although this parameter count is orders of magnitude higher than the number of training samples (680 subjects), the number of parameters we utilize is lower than many of the published state-of-the-art 2D CNNs, despite utilizing 3D convolutions.

### 4. Implementation

All experiments were conducted using python version 2.7.12. The neural network was built with the Keras deep learning library using TensorFlow as backend. TensorFlow, which is developed and supported by Google, is an open-source package for numerical computation with high popularity in the deep learning community. The library allows for easy deployment on multiple graphic processing units (GPUs) (CPU-based experimentation would be prohibitive because of time constraints). The Keras wrapper provides an application programming interface (API) for quicker development and has all functionalities needed to implement the network with the exception of 3D separable convolutions, which we built as a custom layer in TensorFlow. In this paper we employed a Linux machine and two Nvidia Pascal TITAN X graphics cards with 12GB RAM each. The model was parallelized across GPUs such that the feature extractor network works on the AD vs HC and MCI-to-AD conversion problems simultaneously to speed up training. Iterating over the whole training set once, i.e. a single epoch, takes about 30 sec and prediction for a single MCI patient requires milliseconds. Since prediction would not require model parallelization or a lengthy training process, a pre-trained network is practical to be applied on a lower-end GPU (or possibly a CPU) relatively cheaply in a realistic scenario. Across all experiments certain network settings remain unchanged. These include the dropout rate - set at 0.1 for all layers and blocks; the L2 regularization penalty coefficient set at 5*10-5 for all parameters in convolutional and fully connected layers; and the convolutional kernel weight initialization which follows the procedure described by He et al. 2015. The objective function loss is minimized using the Adam optimizer by Kingma and Ba, 2014 with an exponentially decaying learning rate:

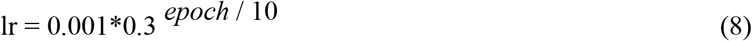

All other parameters are kept at their default value provided in the original Adam paper (Kingma and Ba, 2014). The network hyperparameters were picked because they resulted in sufficiently good performance on the validation set. A training batch size of 6 samples for both the AD and MCI conversion problems is randomly sampled from the dataset when training the network until the dataset is exhausted.

### 5. Performance Evaluation

For the evaluation of the classifier, we repeated the sampling strategy to divide the samples in training, validation and test set splits. Since we have 32 samples more in the MCI dataset (16 for pMCI and 16 for sMCI) as compared to the AD/HC dataset, we used these 32 MCI subjects for testing purposes by randomly sampling 16 subjects from the pMCI and sMCI groups. The validation set comprised roughly 10% of the remaining dataset (36 subjects from MCI and AD/HC respectively) and was also generated by randomly picking in a balanced manner both from the progressive and stable MCI groups and from the healthy and AD patients as we were performing joint learning. Finally, the remaining 340 subjects from both the AD/HC and MCI subsets respectively (i.e. a total of 680 subjects) comprised the training set. No data augmentation procedures were used in this paper.

The model is trained for 40 epochs and the best performing model with the lowest objective function value (eq. 2) on the validation set is saved and its performance is evaluated on the test set. This procedure is then repeated 10 times with different sampling seeds so as to have different samples in the train/validation/test splits (or folds) and minimize the effect of random variation. The number of subjects in each of the training/validation/testing splits in maintained the same at 680/72/32 subjects overall. The trained model is then evaluated on the independent test set. The evaluation metrics used and reported in our results are accuracy (ACC), sensitivity (SEN), specificity (SPE). We also perform receiver operating characteristics (ROC) analysis and compute the AUC across folds. The optimal operating point of the ROC curve was found via Youden’s J statistic. All accuracy, sensitivity and specificity results are reported at the optimal operating point of the ROC curve. For the AD vs HC task, we report the validation results as we only defined a test set for the pMCI/sMCI classification problem (while the AD/HC task is a helpful auxiliary problem, it turned out to be an extremely easy classification problem which is not the focus of this paper).

**Fig. 5.**
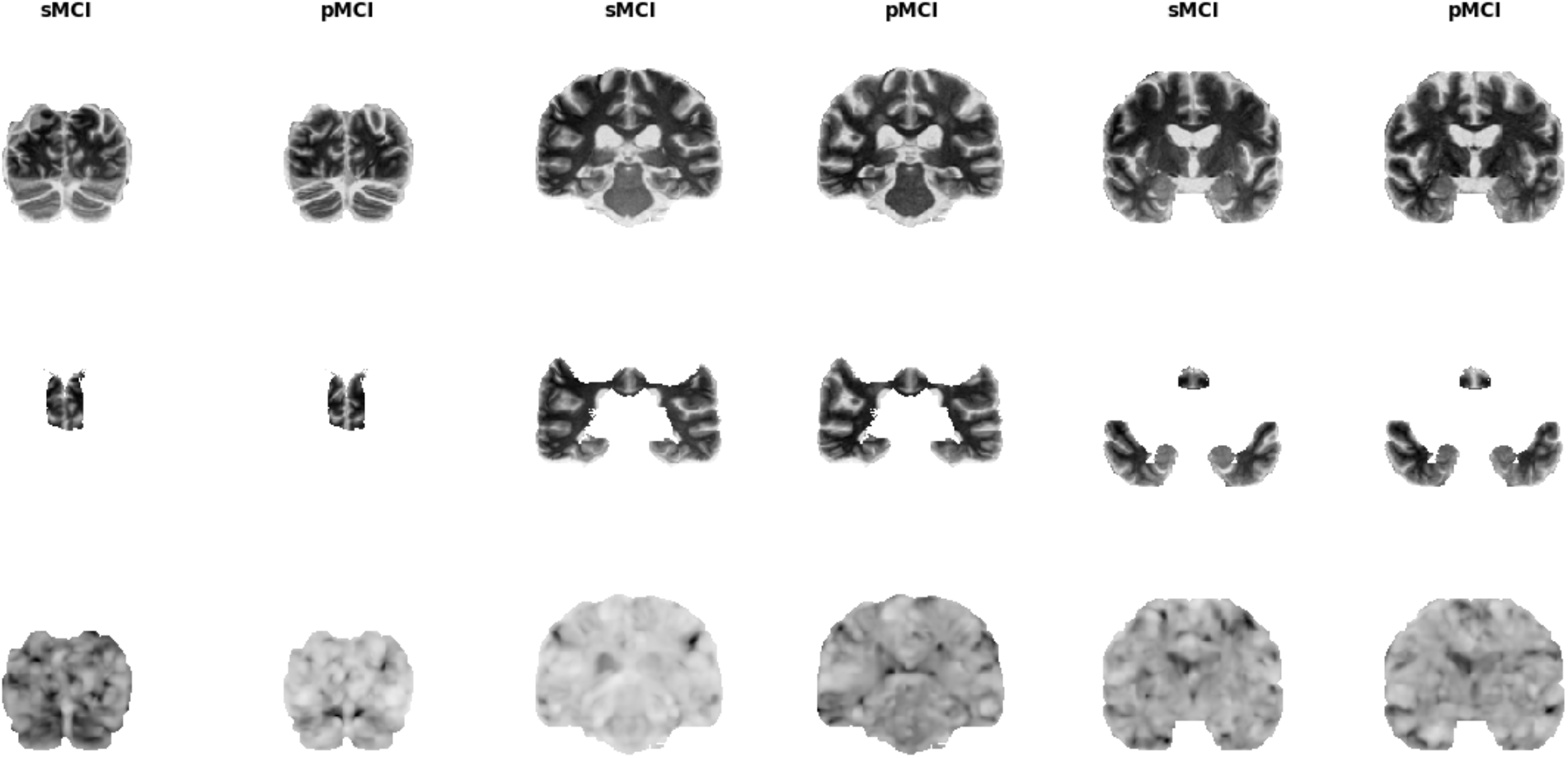
Examples of the image inputs we employ in the classification framework for three different image slices. The upper row shows structural MRI images co-registered to a custom common space. The middle row displays only the brain regions we retain in the atlas-masked tests (parietal, temporal and frontal lobes). The third row shows the Jacobian Determinant images - they indicate the volumetric change a voxel in an unnormalised MRI image must undergo so as to conform to the common template.

### 6. Results

Firstly, we consider the classification performance of our network on four different input biomarker combinations. The four input combinations are: 1) clinical features and T1w MRI images; 2) clinical features and JD images; 3) clinical features and atlas-masked T1w images; and 4) clinical features, JD and T1w images. We performed all of these experiments in our custom template. In order to assess the robustness of the neural network model to MRI structural misalignment, we also performed three experiments in the MNI152_T1_1mm template with three different input combinations (we used all input variants except for 3) clinical features and atlas-masked T1w images). In addition, we assess the performance of our model on the AD vs healthy task with the same input variables as in the pMCI/sMCI problem. We have only included the custom template analyses in the Results section, whereas the MNI space and AD vs healthy experiments are only briefly discussed. A more comprehensive overview on MNI template and AD/HC results can be found in the Supplementary Material. All of our results are based on baseline data from ADNI (no longitudinal data is used in this study).

#### 6.1. Multi-modal classification

Results are summarized in fig. 6 and fig. 7 and table 2.

**Fig. 6.**
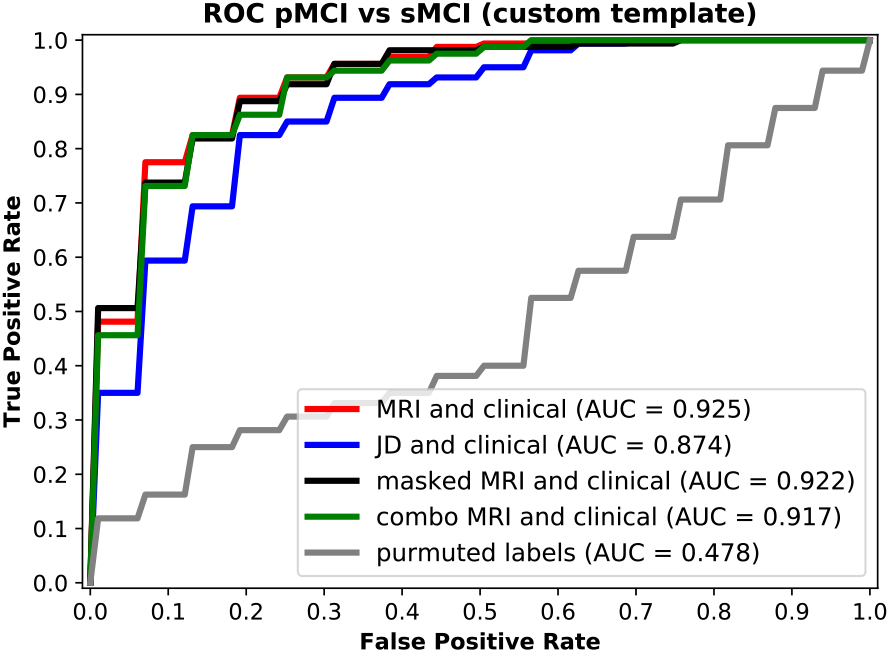
ROC curves of pMCI vs sMCI classification for four input combinations: MRI images and clinical features; JD images and clinical features; Atlas-masked MRI (or just masked MRI) images and clinical features and finally a MRI; and Jacobian Determinant images and clinical features. The MRI data was co-registered to our custom template prior to performing classification. The grey ROC curve at the diagonal was generated by randomly permuting the training labels for the structural MRI and clinical features input combination and predicting using this random classifier.

**Fig. 7.**
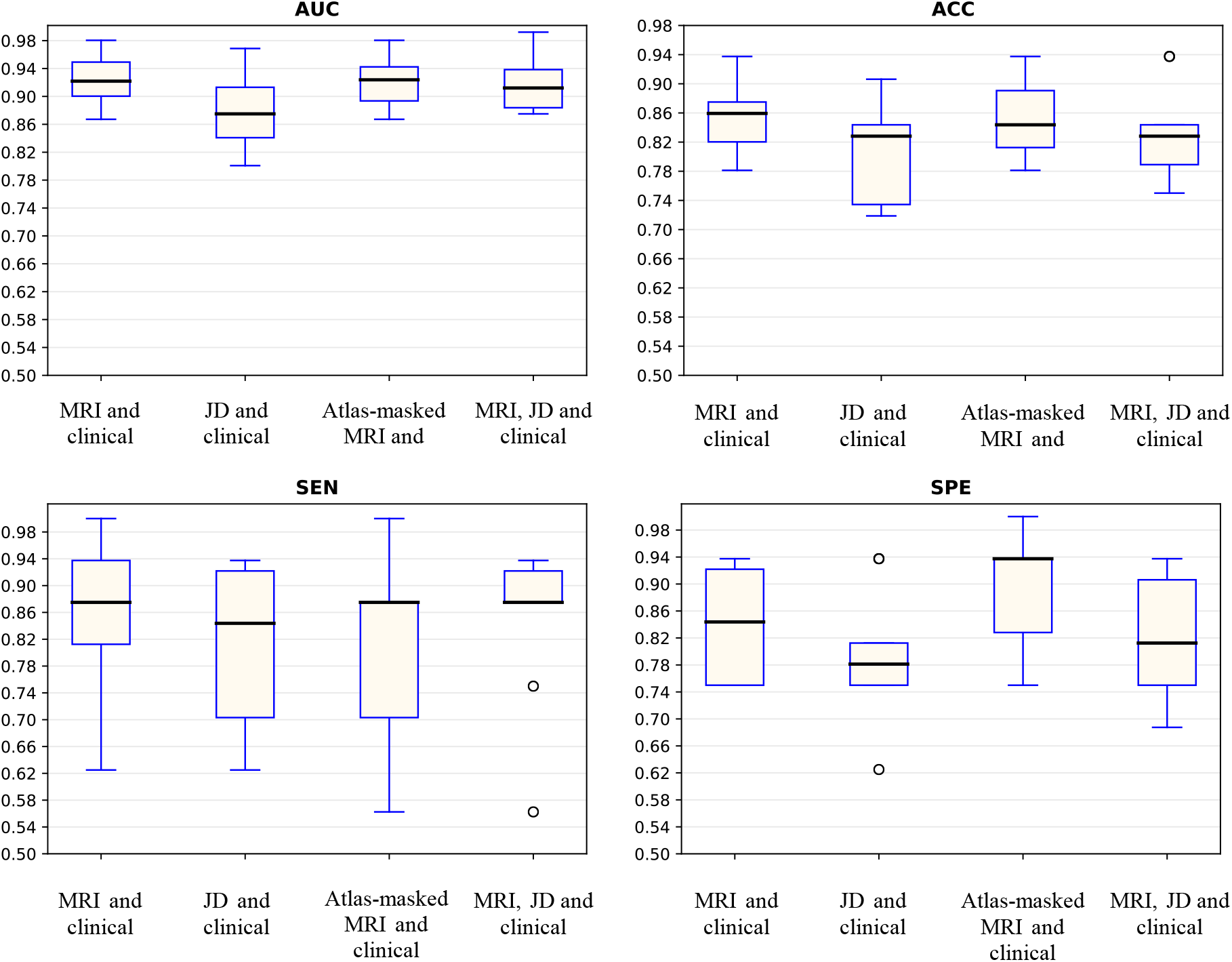
Box plots for AUC, accuracy, sensitivity and specificity for pMCI vs sMCI classification based on multi-stream integration of clinical features and MRI images (co-registered to our custom template) over 10 separate test folds. The black line in each box represents the median value. The boxes encompass values between the 25^th^ and 75^th^ percentile whereas the tails - the top and bottom quartiles. Outliers are marked with a circle. The performance metrics correspond to the optimal operating point of each classifier.

**Table 2.** A comparison table between the median performance metrics on the pMCI vs sMCI classification task using our neural network model.

| pMCI vs sMCI |  |  |  |  |  |  |  |  |
| --- | --- | --- | --- | --- | --- | --- | --- | --- |
| Input Modalities | Custom template |  |  |  | MNI152 template |  |  |  |
|  | AUC | ACC | SEN | SPE | AUC | ACC | SEN | SPE |
| MRI and clinical | 0.925 | 86% | 87.5% | 84% | 0.917 | 85% | 82% | 87% |
| Atlas-masked MRI and clinical | 0.922 | 84% | 87.5% | 94% | - | - | - | - |
| JD and clinical | 0.874 | 83% | 84% | 78% | 0.881 | 82% | 82% | 81% |
| MRI and JD and clinical | 0.917 | 83% | 87.5% | 81% | 0.899 | 83% | 77% | 88% |

The best performance metrics are achieved by including structural MRI along with all clinical data (demographic, neuropsychological, and APOe4 genotyping features). The median AUC across folds for the input combination comprising structural MRI images and clinical features is 0.925 whereas when we remove brain areas not classically associated with AD (i.e. using the Atlas-masked images we employ in the inclusion test), the median AUC obtained is 0.922. Comparing these results across folds using a Mann-Whitney U test indicated that removing brain structures unrelated to the development of AD does not hinder or aid (P=0.4) discrimination in pMCI and sMCI. The median AUC when using JD images and clinical data was found to be 0.874 (Mann-Whitney test yielded p-value=0.041 and 0.046 when compared to the input combinations comprising structural MRI and clinical data, and atlas-masked structural MRI and clinical data results, respectively). Finally, the input combination comprising all types of input streams - T1w images, JD data and clinical features resulted in an AUC of 0.917. Comparing this with the input variants comprising the structural MRI and clinical features, atlas-masked MRI and clinical features, or JD images and clinical features yielded p-values of 0.36, 0.38 and 0.07 respectively (Mann-Whitney-U test). These results suggest that adding structural MRI to the clinical features yields statistically significant higher performance as opposed to using only JD data as an image input stream. In addition, removing brain areas from structural MRI not classically associated with Alzheimer’s disease did not show statistically different classification results compared to the experiments which retained all information. This suggests our model was not negatively impacted by the inclusion of irrelevant or only partially relevant features.

The highest median classification accuracy we achieved was 86%, which resulted from the experiments with structural MRI and clinical data. The atlas-masked MRI and clinical data variant yielded the second best result with 84% classification accuracy, whereas the JD images and the clinical features gave 83% accuracy. Finally, employing all input features also resulted in an accuracy of 83%. Across the classification results from our four different input combinations the median sensitivity varies between 85%-87.5%, and the median specificity between 78% and 94% (evaluated at the optimal point of each curve across the test folds).

Results from the classification performance on both the custom and the MNI152 template are summarized in table 2. We performed Mann Whitney U tests across folds on the obtained AUCs corresponding to the different input combination pairs (custom template vs MNI template). The purpose of these experiments is to assess the robustness of the methodology to possible structural misalignment in the brain areas across images as the MNI space is more “distant” (as compared to the custom template) from the images under study.

The obtained p-values are 0.28, 0.42 and 0.24 for the structural MRI and clinical features, Jacobian Determinants and clinical features, and the combined inputs respectively. Consequently, no statistically significant difference can be found between the performance of our classifier while operating in the two normalization spaces.

Owing to the simpler nature of AD vs HC discrimination, regardless of the input streams and the co-registration template, results are close to 100% on all performance metrics (summarized in table 4 in the Supplementary Material).

**Table 4.** A comparison table between the median performance metrics on the AD vs healthy classification task using our neural network model.

| AD vs HC |  |  |  |  |  |  |  |  |
| --- | --- | --- | --- | --- | --- | --- | --- | --- |
| Input Modalities | Custom template |  |  |  | MNI152 template |  |  |  |
|  | AUC | ACC | SEN | SPE | AUC | ACC | SEN | SPE |
| MRI and clinical | 1 | 99.5% | 100% | 99% | 1 | 99.5% | 100% | 99% |
| Atlas-masked MRI and clinical | 1 | 99.5% | 100% | 99% | - | - | - | - |
| JD and clinical | 1 | 99% | 99.5% | 99% | 0.99 | 97% | 95% | 99% |
| masked and JD and clinical | 1 | 99.5% | 100% | 99% | 1 | 99% | 99% | 99% |

#### 6.2. Classification variance and overfitting

Although we achieve high median performance on all metrics and on both registration templates, dispersion can be further reduced. Fig. 8 shows the standard deviation of the mean training and validation losses across the 10 test folds of the model utilizing structural MRI and clinical features as inputs, which also achieved the highest classification accuracy.

**Fig. 8.**
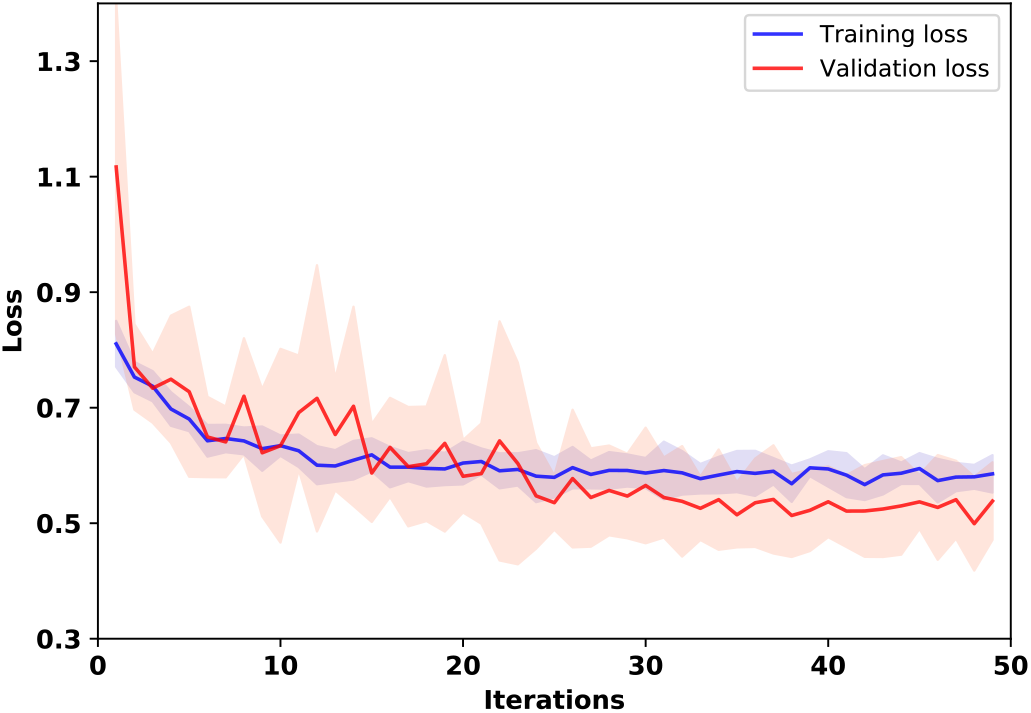
Training and validation losses for our CNN architecture which utilizes structural MRI and clinical features. The standard deviation of the validation loss encompasses the red area in the image, whereas the deviation of the training loss is depicted in blue. The solid lines indicate the means of the losses across the folds.

One factor which contributes to the higher validation variance compared to the training loss curve is the number of samples. Since both the validation and test sets comprise an order of magnitude less subjects than the training set, we also expect the network to manifest higher variance when evaluated on them. Secondly, although the weights were optimized using a variant of stochastic gradient descent, the hyper parameters, such as the dropout rate, the L2 regularization hyper parameter, the initial learning rate and learning rate decay were set to pre-defined values which gave good performance on only one of the validation folds. This was done for two reasons: 1) performing hyper parameter search at each fold was deemed prohibitive given the number of experiments we performed, and 2) it is questionable whether hyper parameter search at each fold would yield clinically relevant results since this cannot be replicated in an applied clinical setting, which would require a predetermined set of hyperparameters. As the dataset is relatively small, we observed some level of overfitting or bias, depending on the specific data split employed. High performance metric variance is most prevalent in the sensitivity and specificity box plots since they are calculated only using either the true positives or true negatives, i.e. half the test set. Accordingly, some studies (Moradi et al. 2015, Hojjati et al. 2017, Tong et al. 2017) repeat their cross-validation loops many times (such as 100 or a 1000 times) in order to further reduce their performance variance, which was not computationally feasible for our deep learning framework.

We would also like to draw the attention of the reader towards the high overlap in the standard deviation between the training and validation losses depicted in fig. 8, indicating comparable performance during both training and validation. Hence, we are confident our network does not suffer from significant overfitting (or underfitting) issues.

### 7. Discussion

Deep learning, or deep neural networks, works by extracting a hierarchy of features from the input data via flexible non-linear transformations. These new data representations are learnt such that they maximize an arbitrary performance metric, for example binary cross-entropy. Hence, instead of relying on expert prior knowledge, or other dimensionality reduction algorithms which might result in a non-optimal set of features, deep neural networks use the gradient in the performance metric to directly guide the feature extraction mechanism. This can result in significant improvements in classification results. Additionally, given that the feature representations are built in a multi-layered fashion (where higher level features are derived from lower level ones), articulate and information-rich images and volumes can be dealt with and incorporated easily into the classification process.

**Table 3.** A comparative table of methodologies on the pMCI vs sMCI classification task using the ADNI dataset. We provide a performance comparison table mainly for recent studies achieving classification rates close to the state-of-the-art. The Methods column includes both the feature selection procedure(s) and the classification method.

| Author | Data | AUC | ACC | SEN | SPE | Conversion time | Validation and Testing method | Method |
| --- | --- | --- | --- | --- | --- | --- | --- | --- |
| Spasov et al. (this paper) | structural MRI + cognitive measures + APOe4 + demographics | 0.925 | 86% | 87.5% | 85% | 0-36 months | 10-fold cross-validation | CNN |
| Hojjati et al. 2017 | rs-fMRI | 0.95 | 91.4% | 83.24% | 90.1% | 0-36 months | 9-fold cross-validation (report on validation set) | Graph measures + SVM |
| Moradi et al. 2015 | structural MRI + cognitive measures | 0.9 | 82% | 87% | 74% | 0-36 months | 10-fold cross-validation (report on test set) | LASSO + SVM |
| Liu et al. 2017 | structural MRI + FDG-PET + cognitive measures + APOe4 + demographics | 0.92 | 84.6% | 86.5% | 82.4% | 0-36 months | holdout | ICA + Cox model |
| Korolev et al. 2016 | structural MRI + clinical data + plasma-proteomic data + medications | 0.87 | 80% | 83% | 76% | 0-36 months | 10-fold cross-validation (report on test set) | Joint Mutual Information + Kernel Learning |
| Beheshti et al. 2017 | structural MRI | 75.08 | 75% | 77% | 73% | 0-36 months | 10-fold cross-validation | Morphometry + t-test + SVM |
| Choi et al., 2018 | flurodeoxyglucose and florbetapir PET | 0.89 | 84.2% | 81% | 87% | 0-36 months | holdout | CNN |
| Tong et al., 2017 | structural MRI + cognitive measures | 0.92 | 84% | 88.7% | 76.5% | 0-36 months | 10-fold cross-validation (report on test set) | Elastic Net + SVM |
| Lu et al. 2018a | FDG-PET | - | 82.5% | 81.4% | 83% | 0-36 months | 10-fold cross-validation | NN |

In this paper, we developed a new method with the primary goal of early identification of MCI patients with high risk of converting to Alzheimer’s disease up to three years prior to diagnosis, and the subsidiary task of Alzheimer’s patient vs. healthy control discrimination. Our approach uses a parameter-efficient deep convolutional neural network framework, inspired by grouped and separable convolutions, to extract descriptive factors from structural MRI images acquired at baseline. In this respect our work differs from previous deep learning-based methods for early AD detection in that it takes into consideration data paucity in medical datasets and introduces design precautions by reducing the number of network parameters. This in turn increases the generalization capabilities (i.e. reduces overfitting) of our model to unseen test samples, thus enabling us to achieve state-of-the-art MCI-to-AD classification performance. The structural MRI images are complemented by standard cognitive test results (CDRSB, ADAS, RAVLT), demographic information (age, gender, ethnic and racial categories, education) and APOe4 expression levels also acquired at baseline to arrive at a final score which is used to predict conversion. We chose these biomarkers in order to create a classification methodology which is as minimally invasive as possible. Hence, for example, we do not include PET imaging because of radiation exposure and CSF data owing to the potentially painful lumbar puncture which can also lead to clinical complications. Additionally, we exploited AD/HC data to limit the effects of overfitting. This was achieved by multi-task learning where the same network layers are used to extract representations from the input biomarkers for both the MCI-to-AD conversion task and the AD/HC classification problem. While previous methods employ pre-training (Payan et al. 2015; Hosseini-Asl et al. 2016; Liu et al. 2018) to reap similar benefits, this requires training a model twice, whereas dual-learning is a single-stage procedure, hence facilitating training. Also, we assessed the performance of our method using two different co-registration templates (a custom template and the MNI152 template) as well as various input combinations of structural MRI, the local JD of the deformational field computed during MRI co-registration, as well as the clinical data. The best result we obtained was a mean AUC of 0.925 averaged across 10 different testing folds with a mean MCI-to-AD conversion prediction accuracy of 86%, sensitivity of 87.5% and specificity of 85% (see table 2). It is also important to note that, to the best of our knowledge, the only study which presents better classification results on the pMCI vs sMCI problem (Hojjati et al. 2017) does not explicitly mention the use of separate test set, possibly leading to circular analysis (results are reported on a validation set instead of a dedicated test set).

The main novelties of our method were 1) the use of parameter-efficient layers, such as grouped and separable convolutions (implemented as custom Keras layers for 3D inputs) which reduce the number of network parameters, hence limiting overfitting; 2) the substitution of network pre-training, which was typical in earlier deep-learning based AD classification studies (Payan et al. 2015, Hosseini-Asl et al. 2016), with multi-task learning which utilizes AD/HC data to arrive at a single-stage training approach and 3) the utilization of the JD as a complementary imaging input stream to maximize the extracted information from the structural MRI.

Convolutional neural networks abstract away the manual handcrafting of useful features from medical images, such as the use of pre-defined brain regions of interest (Da et al. 2013). Intuitively, neural network- based methods should perform better as the feature extraction process is directly driven by the performance optimization procedure, however, it comes at the cost of a relatively high number of network parameters compared to the number of samples. Since there are no formal estimates of the number of training samples required for a given convolutional architecture to achieve good generalization, we are driven by the metaheuristic approach of minimizing the number of network weights and maximizing the effective number of training examples so as to boost performance on an independent test set and consequently during clinical application. As a result, our 3D model comprises ~550,000 parameters, which is orders of magnitude lower than conventional 3D CNNs and even lower than recent 2D CNNs, such as AlexNet (Krizhevsky et al., 2012) and Xception (Chollet, 2017). This was not done by sacrificing network depth or structural complexity but rather by inserting efficient convolutional layers. In order to facilitate the learning procedure, we hypothesized that employing an auxiliary task and minimizing the joint training objective of the MCI-to-AD conversion and AD/HC classification tasks would be an effective alternative to pre-training. In this context, AD/NC discrimination is seen as a simpler version of MCI conversion prediction, and in order to speed up training convergence we worked under the assumption that similar descriptive factors would be useful for both problems.

Considering existing computer vision research, deep learning methodologies for computer-aided diagnostics would also be applicable on non-co-registered or even non-pre-processed images, however, this approach could lead to image artefacts contributing to the discriminatory performance of the algorithm, which could learn to relate center-specific (rather than disease-specific) features with disease outcomes. As with all multicentric studies, careful and unified data collection and processing is crucial to minimize this confound.

Comparing our classification metrics with recent studies indicate that only Hojjati et al. 2017 who use rs-fMRI outperform our results (although, as mentioned above, only reporting on a validation set comprising 4 subjects via 9-fold cross-validation). Unfortunately, at the time of writing ADNI provides limited rs-fMRI data (18 pMCI and 62 sMCI subjects) so it would be difficult to predict how their results would scale to larger populations. Additionally, using structural MRI only can significantly reduce in-patient scanner time as opposed to including a functional scan. To the best of our knowledge, the study by Liu et al. 2017 is the first to produce comparable performance (at least in some metrics) to our model, at 84.6% classification accuracy vs 86% for our work. The difference is, however, that Liu et al. 2017 utilize FDG-PET as an extra modality which is known to be extremely informative in AD, as well as structural MRI and all the biomarkers we have employed. Moradi et al. 2015 and Tong et al. 2017 both use a very similar methodology to each other and the same data (structural MRI and cognitive assessment tests) as in this paper. Their sensitivity metrics are comparable to our model at ~ 87%-88% but manifest lower specificity at 74% and 76% respectively, while our deep learning method achieves a median specificity of 85% across folds (and 94% specificity when using the Atlas-masked MRI and clinical features as inputs). A possible explanation would be the inclusion of APOe4 and demographic data as well as the efficacy of the neural network. Also, as is discussed in Moradi et al. 2015 the labelling and number of ADNI subjects varies across studies, thus hampering direct comparisons.

In summary, we developed a deep learning-based method for the early prediction of MCI-to-AD converts by combining structural MRI, neuropsychological assessment data and APOe4 expression levels obtained from the ADNI database at baseline. We achieved a very high predictive performance with an average AUC of 0.925, prediction accuracy of 86%, sensitivity of 87.5% and specificity of 85%. Our study proposes the use of more efficient neural network architectures comprising fewer parameters to limit the effects of overfitting. The convolutional framework is generic and applicable to any 3D image dataset and gives the flexibility to design a computer-aided diagnosis system targeting the prediction of any medical condition utilizing multi-modal imaging and tabular clinical data.

## 8. Acknowledgements

In this work we employed the database of the Alzheimer’s Disease Neuroimaging Initiative (ADNI). ADNI was formed as a multicenter longitudinal study to identify imaging, clinical, genetic and biochemical biomarkers for the early detection and tracking of Alzheimer’s disease (AD) and Mild Cognitive Impairment (MCI). ADNI is the result of a $67 million partnership by the public and private sector. Financial support was obtained from the National Institute on Ageing, 13 pharmaceutical companies, and two foundations providing funding through the Foundation for the National Institutes of Health. The study can be split in three sub-initiatives - ADNI1, ADNI2 and ADNI GO. The initial phase known as ADNI1 included subjects between 55-90 years of age from approximately 50 sites from the US and Canada. ADNI2 and ADNI GO add new participants and funding to the study. The database is made available to researchers around the world and has a broad range of collaborators. The principle investigator of ADNI, who oversees all aspects, is Dr. Michael Weiner, MD, VA Medical Center and University of California - San Francisco. For up-to-date information, see www.adni-info.org. Simeon Spasov is supported by the Engineering and Physical Sciences Research Council [EP/ L015889/1]. Luca Passamonti is funded by the Medical Research Council (MRC) grant (MR/P01271X/1) at the University of Cambridge, UK.

## Supplementary Material

**Fig. 9.**
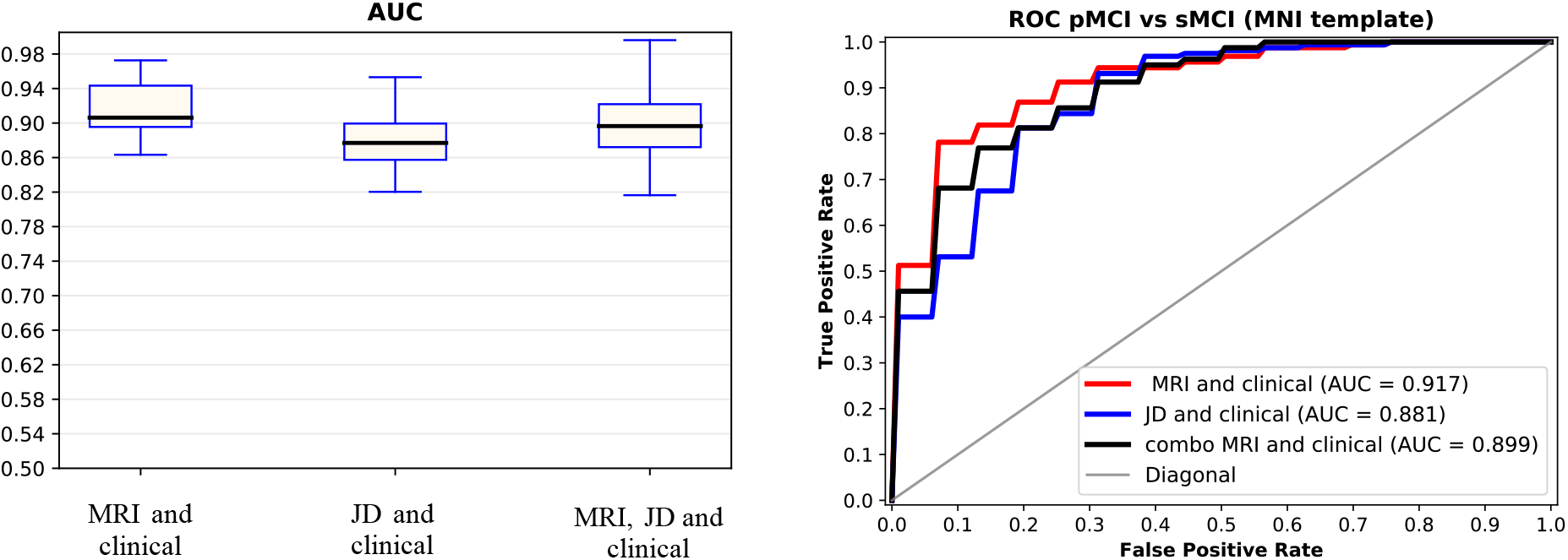
Box plots for AUC, accuracy, sensitivity and specificity obtained on the pMCI vs sMCI classification task from structural MRI, Jacobian Determinant and atlas-masked structural MRI inputs (all using clinical features) over 10 separate test folds. The MRI data is coregistered to the MNI(152) template. The black line in each box represents the median value. The boxes encompass values between the 25^th^ and 75^th^ percentile whereas the tails - the top and bottom quartiles. Outliers are marked with a circle.

